# Landing the ‘Tiger of Rivers’: Understanding Recreational Angling of Mahseers in India using YouTube Videos

**DOI:** 10.1101/2023.07.22.550129

**Authors:** Prantik Das, V.V. Binoy

**Affiliations:** The University of Trans-Disciplinary Health Sciences and Technology (TDU), Bangalore, India; National Institute of Advanced Studies (NIAS), Indian Institute of Science (IISc) Campus, Bangalore, India

**Keywords:** Mahseer, Recreational Angling, Anglers, Human-Animal Interactions, YouTube, Digital Data, Conservation Culturomics, iEcology.

## Abstract

Megafish mahseers popularly known as the ‘tiger of rivers’, are the dream catch of recreational anglers in India. The present study explored the Recreational Angling (RA) videos of five mahseer species *Tor khudree* (deccan mahseer), *T. putitora* (golden mahseer), *T. remadevii* (humpback mahseer), *T. mosal* (mosal mahseer) and *Neolissochilus hexagonolepis* (chocolate mahseer) recorded from India and uploaded on the social media platform YouTube from January 2010 to October 2022. We did not come across any RA videos of *T. mosal* and *T. remadevii* on YouTube hence further analyses were carried out on the remaining three focal species. No seasonality was observed in the frequency of RA videos uploaded on YouTube and *T. khudree* attracted the highest number of views per video. Catch and Release (C&R), an ethical RA practice was noticeably low in the case of *N. hexagonolepis*. The size of the catch was found to be positively associated with the social engagement received by the RA videos of all the three mahseer species focused. Angler and angling-related remarks and words associated with the emotion ‘trust’ dominated the comments received by the videos. The results are discussed in light of the trending discourses on developing social media data as a complementary tool for monitoring and managing RA and conserving fish.

## Introduction

The Mahseers are a group of freshwater megafishes, belonging to three genera, viz. *Tor, Neolissochilus,* and *Nazitor,* of the Cyprinidae family. Popularly known as the ‘Tiger of Rivers’ these iconic fishes are distributed across the rivers of South and South-East Asia (Pinder and Raghavan 2013; Nautiyal 2014). The large body size requiring a substantial quantity of biologically significant resources for survival and reproduction makes these megafishes vulnerable to the modifications happening in their habitat. Natural populations of mahseers in many nations are threatened by human activities like river flow alteration and regulation (construction of dams, hydropower projects, canals etc.), water pollution, alteration of drainage basins (deforestation), over-harvesting (Dudgeon 2000; Raghavan et al. 2011; Bhatt and Pandit 2016; Everad et al. 2018), etc. Amongst various anthropogenic threats that different mahseer species face, Recreational Angling (RA) - “fishing of aquatic animals (mainly fish) that do not constitute the individual’s primary resource to meet basic nutritional needs and are not generally sold or otherwise traded on export, domestic or black markets” (FAO 2012) - is a matter of great concern.

Experiencing and overcoming the challenges posed by the fish caught on the hook is the main motivator for the recreational anglers (Fedler and Ditton, 1994; Beardmore et al. 2011; Young et al. 2016). Many anglers follow Catch and Release (C&R), where the captured fish is unhooked and released alive into the water after measuring its length and weight and clicking photographs. Angling has been a size selective activity (Arlinghaus and Mehner 2003; Isermann et al. 2005) and the goal of each angler is to catch the biggest fish called the ‘prize or trophy catch’ hence achieving the pride and satisfaction (Everard and Kataria 2011). Although RA does boost the local economy (Hyder et al. 2018; Arlinghaus et al. 2019) by promoting tourism, this activity also causes an extremely high number of fishes being harvested across the world (Cooke and Cowx 2004; Hyder et al. 2018; Brownscombe et al. 2019; Sbragaglia et al. 2020). Furthermore, the stress and injury happening to the individuals from constant hooking, unhooking and physical exertion could affect the health of the exploited population eventually causing a decline in their fitness (Cooke and Cowx 2006; Arlinghaus et al. 2007a; b; Cooke and Sneddon 2007). Improper hooking locations (deep hooking in gills, oesophagus, etc.), injuries caused by the gear, and inappropriate handling practices such as prolonged duration of fight and air-exposure during and after the catch by amateur and inexperienced anglers can even lead to the death of the angled fish (Cooke and Cowx 2004; Meka and McCormick 2005; Cooke and Cowx 2006; Lewin et al. 2019). Truncation of natural age and size, loss of genetic variability and trophic level changes are also reported in the fish populations exploited for RA (Schmitz et al. 2004; Cooke and Sneddon 2007). Arlinghaus et al. (2017a) argue that angling can result in ‘gear-induced timidity syndrome’- the transformation of the exploited population into consistently more timid/shy in comparison to their unexploited counterparts. Modifications in the behaviour, social interaction patterns, life history, food web dynamics (Arlinghaus et al. 2017a) induced by the timidity syndrome could have severe consequences at the levels of population and ecosystem. Other issues related to RA such as the episodes of losing or leaving fishing gear in the water, littering of nylon lines, poisoning caused by the lost angling gears (sinkers, jigs) containing heavy metal lead (Scheuhammer et al. 2003) etc. could deteriorate the health of the water bodies (Cowx 2002; Asoh et al. 2004).

Along with their enormous size, mahseers are also known for their resistance and fighting power that makes them one of the greatest and the most popular game fishes in the Indian subcontinent (Thomas 1873; Dhu 1906; 1923; MacDonald 1948). India accounts for about 16 of the 47 mahseer species found in the world (WWF 2013; Pinder et al. 2019) and *T. khudree, T. putitora, T. remadevii and N. hexagonolepis* have been dominantly angled across India since the early 19^th^ Century (Gupta et al. 2015b; Bower et al. 2017). The rush and thrill that the mahseer provides to the anglers from the great fight that they put up during angling is one of the reasons why professional recreational anglers from all over the globe visit India to angle this much prized fish (Kulkarni and Ogale 1978; Gupta et al. 2015a; Bower et al. 2017). Furthermore in India, C&R, an ethical angling practice, had been promoted from the mid-1970s as a strategy to protect the wild mahseer populations from the illegal fishing activities and poachers (Everard and Kataria 2011; Pinder and Raghavan 2013; Baruah and Sarma 2018). However in the year 2012 Supreme Court of India equated C&R angling to baited hunting under the Indian Wildlife Protection Act (WPA 1972) and subsequently banned this activity in all the protected areas (PA) of the country (Ajay Dubey vs. National Tiger Conservation Authority (NTCA): special leave petition no(s).21339/2011; Gupta et al. 2015b)

Although understanding the pressure of RA that mahseer populations face in different rivers of India is imperative to design effective strategies for their conservation, no official guidelines for monitoring and managing RA is available in India till date (Gupta et al 2015a). Monitoring the angling activities happening at numerous rivers of a geographically large and culturally diverse nation like India and estimating its impact on different mahseer species is an activity requiring enormously huge financial, human and technical resources (Arlinghaus et al. 2017b; Brownscombe et al. 2019).

Advent of the digital era and the popularisation of social media not only generated a huge amount of data on every aspect of nature and society but also witnessed the arrival of new branches of science, ‘iEcology’ – the study of the patterns and processes of ecology using the data available on the internet (Jarić et al. 2020) and ‘conservation culturomics’ - the study of human-nature interactions for understanding conservation issues using digital datasets generated by people on the internet (Ladle et al. 2016; Correia et al. 2021). The digital data produced and uploaded on a variety of platforms by people from various cross sections of the society could function as a representation of their ideas, beliefs, concerns and emotions towards the living and nonliving components of the ecosystem (Hamilton and Carlston 2013; Correia et al. 2016) and may even reflect the nature of the human-animal relationship existing in their community (Lennox et al. 2020; Herrera et al. 2023). Using the internet and social media data as a complement to the extremely resource demanding conventional methods popularly used for monitoring human-animal interactions including RA, has become a trend in the recent past (Carter et al. 2015; Venturelli et al. 2017). For instance, many researchers have used citizen science based social media data to explore the impact of marine RA in the Mediterranean (Giovos et al. 2018; Sbragaglia et al. 2020; 2021; Eryasar and Saygu 2022) and in West-African countries (Belhabib et al. 2016). Unfortunately, such an attempt to use the internet/social media data to get insights on the impacts of RA on mahseers has not commenced yet.

With the availability of pocket friendly and easily accessible videography and editing tools, YouTube a video-sharing website has evolved as a very popular medium of self-expression and communication (Burgess and Green 2018). An exploration of this platform reveals that many recreational anglers upload videos of their angling and the catch making it a valuable digital resource for the researchers interested in RA (Struthers et al. 2015; Gioves et al. 2018; Sbragaglia et al. 2020; Eryasar and Saygu 2022). The attention that anglers get on the digital space can work as social rewards as well as the basis for financial benefits given by the platform. Hence, YouTube works not only as a repository, but also offers a platform for bidirectional communication between the anglers and the general public. The views, likes and comments with emoticons added by the spectators on the videos could have an impact on the content of the future videos created and posted by the anglers (Sbragaglia et al. 2020; Lennox et al. 2022). Hence, along with giving insights on the behaviour of anglers in the natural fish habitats, YouTube videos can also provide valuable information on the responses of the society towards the practices followed by the anglers.

The present study explored the following questions focusing the angling videos of five mahseer species found across India, viz. *T. khudree* (Deccan mahseer/blue-finned mahseer), *T. remadevii* (humpback/orange-finned mahseer), *T. putitora* (golden mahseer), *T. mosal* (mosal mahseer) and *N. hexagonolepis* (chocolate mahseer) available on YouTube:

1. Is there any seasonality in the frequency of the angling videos featuring the focal mahseer species recorded from different states of India uploaded on YouTube?
2. Do anglers meticulously follow C&R for each of the focal species?
3. Do the species and the size (Total Length; TL) of the catch decide the Social Engagement (SE; the number of views, likes, and comments) and sentiment of the comments received by the mahseer angling videos?

## Materials and methods

The angling videos of five different mahseer species reported from different regions of India, *T. khudree, T. remadevii, T. mosal, T. putitora* and *N. hexagonolepis* uploaded on YouTube from January 2010 to October 2022 were extracted using the keywords given in SM1. These videos were manually sorted based on the name of the state, river and/or region mentioned in the title or the description. The videos that did not show mahseer getting caught or those that were recorded from locations other than India were not considered for the analysis. However, the comments received by the former were subjected to sentiment analysis to know the nature of the response they received from the viewers. From each video the following information was extracted: month and year of upload, region/state where angling was carried out, size (Total Length; TL) of the catch, number of views, likes and comments, the content of the comments and whether C&R was practised. The size of the fish (TL) was obtained from videos or descriptions available with it. In cases where no such information was available, TL of the individual caught was determined by keeping the details of the lures available on their company website (Rapala J13, Mepps Spinner, Spoon lure) as the reference. Angled fish not released back, taken afar from the shore, placed inside a bag, killed or cooked, were considered as cases where C&R was not being practised. While analysing the videos, we found Invasive Alien Fishes (IAF) being caught accidentally by some of the anglers targeting mahseers. Since IAF are emerging as a threat to indigenous species including mahseers (Raghavan et al 2011; Gupta et al 2020) and the exploration of internet data is evolving as a tool to get insights on their presence and distribution in an ecosystem (Mamun et al. 2023) each video collected was checked for the presence of such species. Mining of YouTube data, available on the public domain, does not require any specific ethical clearance. However, in order to maintain the anonymity of the people who uploaded the videos on YouTube, no details that could be used to identify the individuals like personal information, channel id, channel title, etc., were collected.

## Data analysis and statistics

In order to study the annual rhythms (called seasonality; Sbragaglia et al. 2020) in the upload patterns of the mahseer angling videos on YouTube, RAIN (Rhythmicity Analysis Incorporating Nonparametric methods) was conducted on RStudio version 4.1.2. using ‘BiocManager’ and ‘rain’ packages (Thaben and Westermark 2014). We used Generalised Linear Modelling (GLM) with the Poisson family distribution for the analysis as the data was not normally distributed. The size of the catch (TL) was taken as the dependent variable while the SE received by the videos (number of views, likes and comments) and the sentiment scores of comments were the independent variables. For C&R, a score of 1 was given if fish were released and 0 for not practising this ethical angling protocol. In order to understand the species specific variation in the SE, Kruskal-Wallis Rank Sum Test followed by a *post-hoc* analysis (Dunn Test with Bonferroni method) was conducted.

The comments received by the angling videos were classified into six categories: fish related, angler related, angling technique related, location related, IAF related and miscellaneous. The last category included greetings, spam comments, non-contextual mentions and links, etc. The repeated comments were also removed. All categories barring the ‘miscellaneous’ were further processed; the text was lemmatized by removing the special characters, numbers, punctuations, English stop-words such as “is”, “the”, “and”, “a”, etc., and white spaces using the ‘tm’ package on RStudio version 4.1.2. and was subjected to the following analysis:

a. *Word Frequency Analysis*: in which the top five keywords that occurred the maximum number of times in the comments of the videos were recorded.

b. *Sentiment and Emotion Classification Analysis*: sentiment scores were given as positive, neutral and negative on an integer scale of -5 to +5 using the AFINN Lexicon (Nielsen 2011). Emotion classification was built on the NRC Word-Emotion Association Lexicon (EmoLex) consisting of a list of English words and their associations with 8 basic emotions viz. anger, fear, anticipation, trust, surprise, sadness, joy, and disgust (Mohammad and Turney 2010; 2011).

All analyses were done on R version 3.6.1. and RStudio version 4.1.2. using ‘FSA’, ‘wordcloud’, ‘syuzhet’ ‘RColorBrewer’ and ‘ggplot2’ packages.

## Results

A total of 108 angling videos of different mahseer species, which accounted for 245 individual fish, were obtained from YouTube. However, 15 videos without a mahseer getting caught, appearing in the results due to the presence of keywords used for the search in their title or the description, had to be excluded. The remaining 93 videos (16.1% - *T. khudree* 40.9% - *T. putitora* and 43% - *N. hexagonolepis*) were considered for the analyses. Interestingly, we did not get any videos of *T. mosal* and *T. remadevii* being angled. The IAF tilapia (*Oreochromis spp.*) was found hooked in three *T. khudree* angling videos. The angling sites of these videos with IAF were in the states of Maharashtra and Kerala. The other bycatch species found in 4 of the 108 angling videos were *Labeo rohita* (rohu), *Bagarius yarrelli* (goonch) and *Schizothorax richardsonii* (common snowtrout).

Only a handful of angling videos of *T. putitora* and *T. khudree* recorded from various states of India were uploaded on YouTube during the time period 2010 to 2012. For the next three years no videos of these two species were added to this platform (Fig. 1). However, from the year 2015 onwards more videos depicting these two species appeared on YouTube. Videos of *N. hexagonolepis* uploaded before the year 2017 were not available on YouTube, while the number peaked in 2022. No significant annual oscillations (seasonality) were observed in the angling videos uploaded on YouTube across the three focal mahseer species (p > 0.05; Fig. 2). Major angling sites of the *T. khudree* videos were in the state of Karnataka (60%) while *T. putitora* and *N. hexagonolepis* were mainly caught from Uttarakhand (37%) and the North-Eastern state of Sikkim (50%; Fig. 3) respectively. Interestingly, none of the angling activities in the videos were undertaken in areas outside the geographical distribution of the respective species. Unfortunately, only 39% of *N. hexagonolepis* individuals were released into the river after being caught by the anglers (30% of videos). In contrast, this ethical angling practice (C&R) was noted with 59.1% of *T. khudree* (60% of videos) and 47.4% of *T. putitora* individuals (57.9% of videos; Fig. 4).

**Figure 1.**
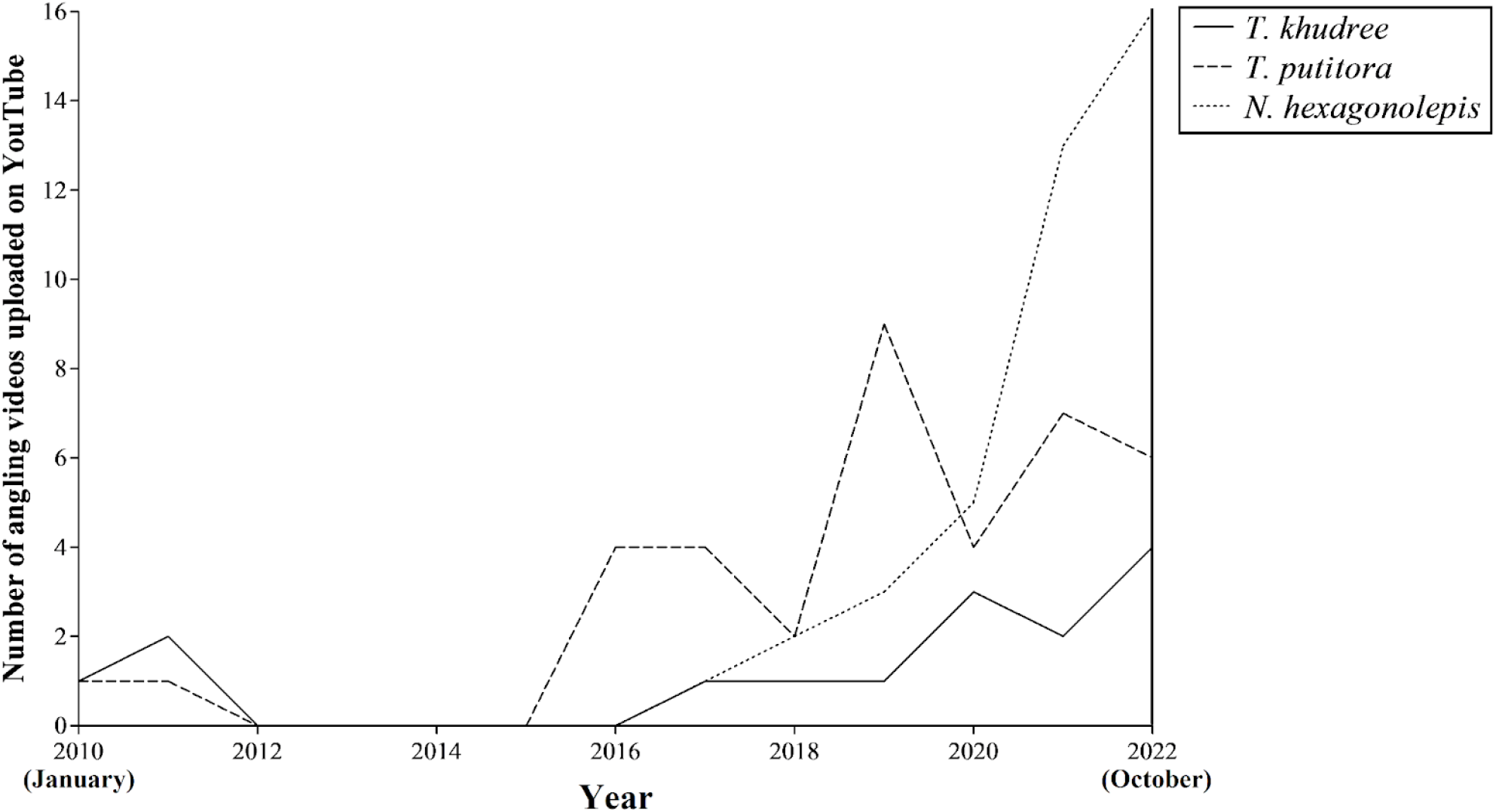
The number of angling videos of the three focal mahseer species uploaded on YouTube from January 2010 - October 2022.

**Figure 2.**
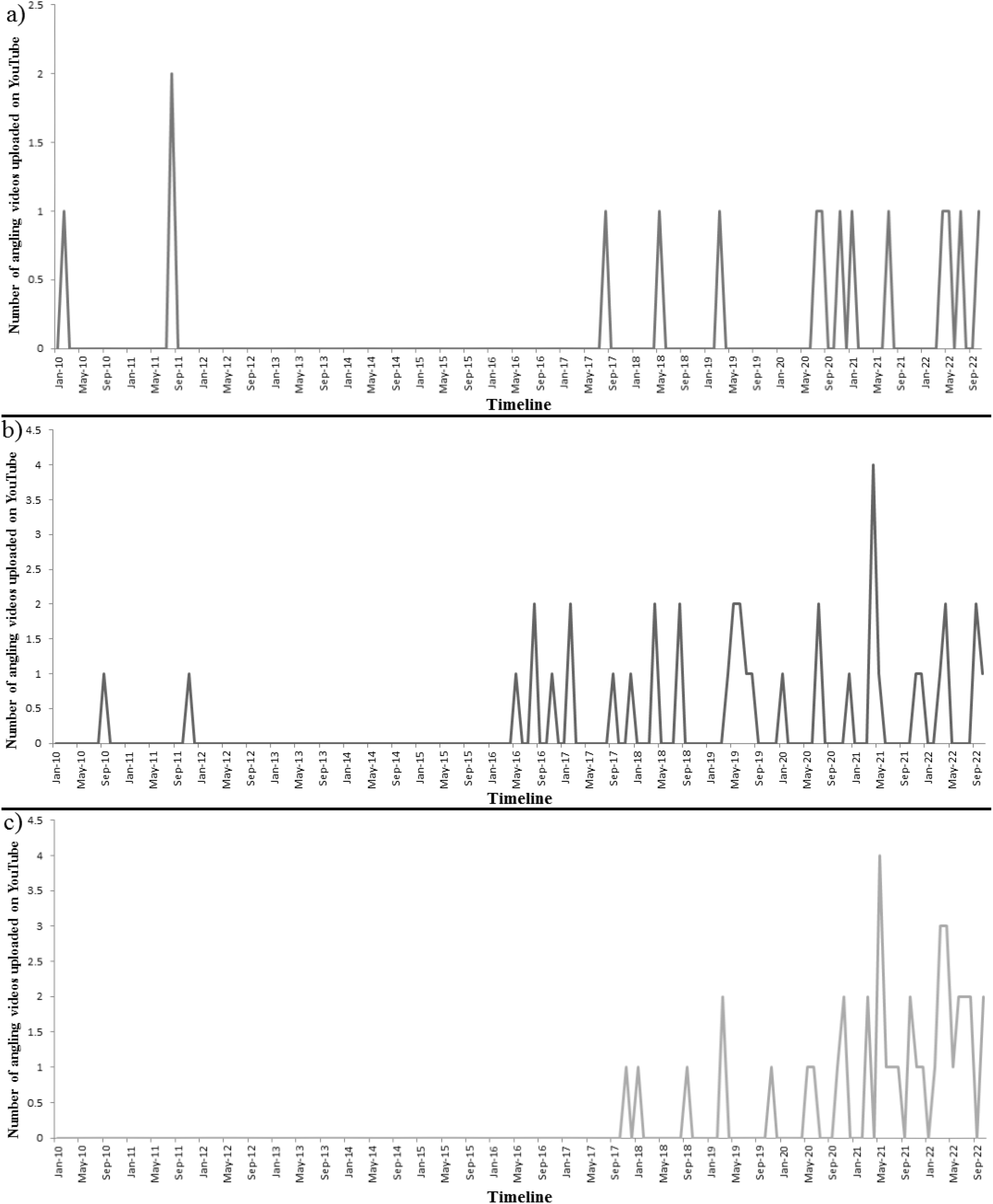
Monthly upload of the angling videos of a) *T. khudree*, b) *T. putitora* and c) *N. hexagonolepis* on YouTube (from January 2010 to October 2022).

**Figure 3.**
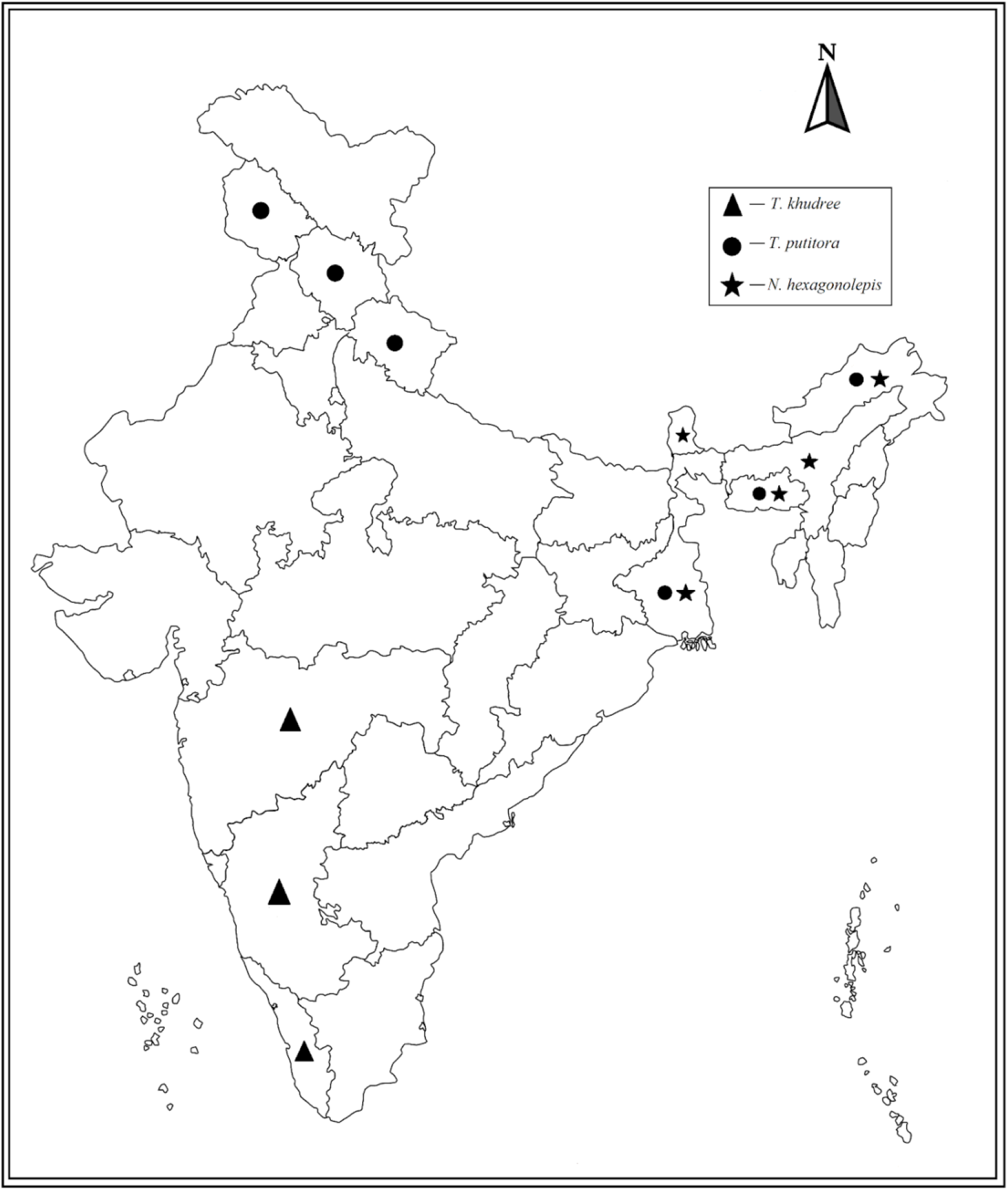
Map of India displaying the states from where the angling videos of the three focal mahseer species were recorded. Map not to scale.

**Figure 4.**
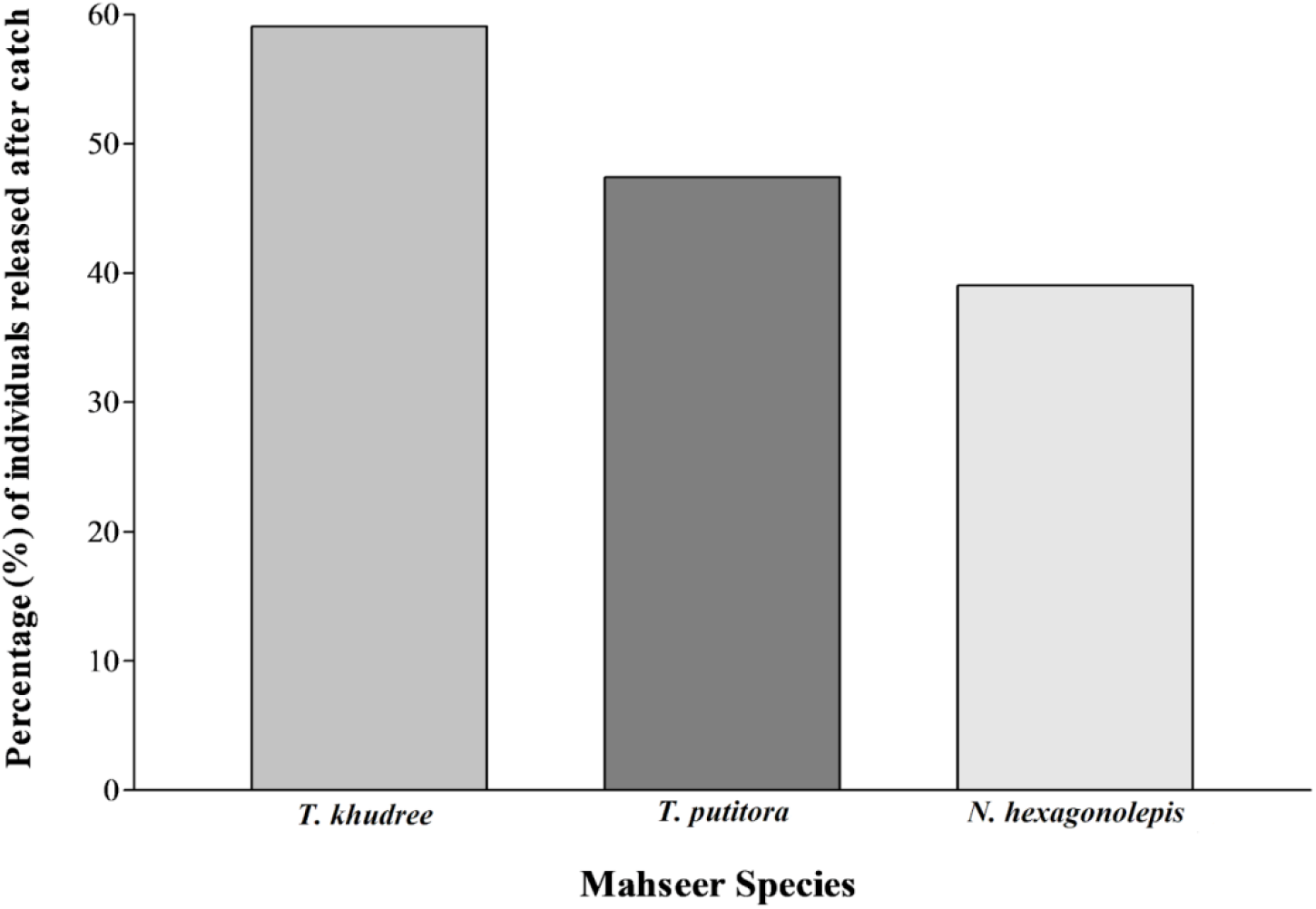
Percentage (%) of individuals of the three focal mahseer species released after catch in the angling videos.

We found a significant difference in the number of views per video across the three focal species (Kruskal-Walllis Rank Sum Test; χ^2^ = 8.62, df = 2, p = 0.01). A *post-hoc* analysis revealed that videos of *T. khudree* received a significantly higher number of views in comparison to the *N. hexagonolepis* (Dunn Test; Z = -2.91, p = 0.01) and *T. putitora* (Dunn Test; Z = 2.34, p = 0.04; Fig. 5). No interspecies difference was observed in the number of likes (χ^2^ = 5.40, df = 2, p = 0.06) and comments received per videos (χ^2^ = 5.65, df = 2, p = 0.06). All elements of SE; number of views, likes, comments were found to be positively correlated with the catch size (TL) in all three focal species of mahseer (Fig. 6); *T. khudree* (views β = 0.12, p < 0.001; likes β = 0.08, p < 0.001; comments β = 0.02, p < 0.001), *T. putitora* (views β = 0.02, p < 0.001; likes β = 0.01, p < 0.00; comments β = 0.01, p < 0.001) and *N. hexagonolepis* (views β = 0.03, p < 0.001; likes β = 0.01, p < 0.001; comments β = 0.005, p < 0.01).

**Figure 5.**
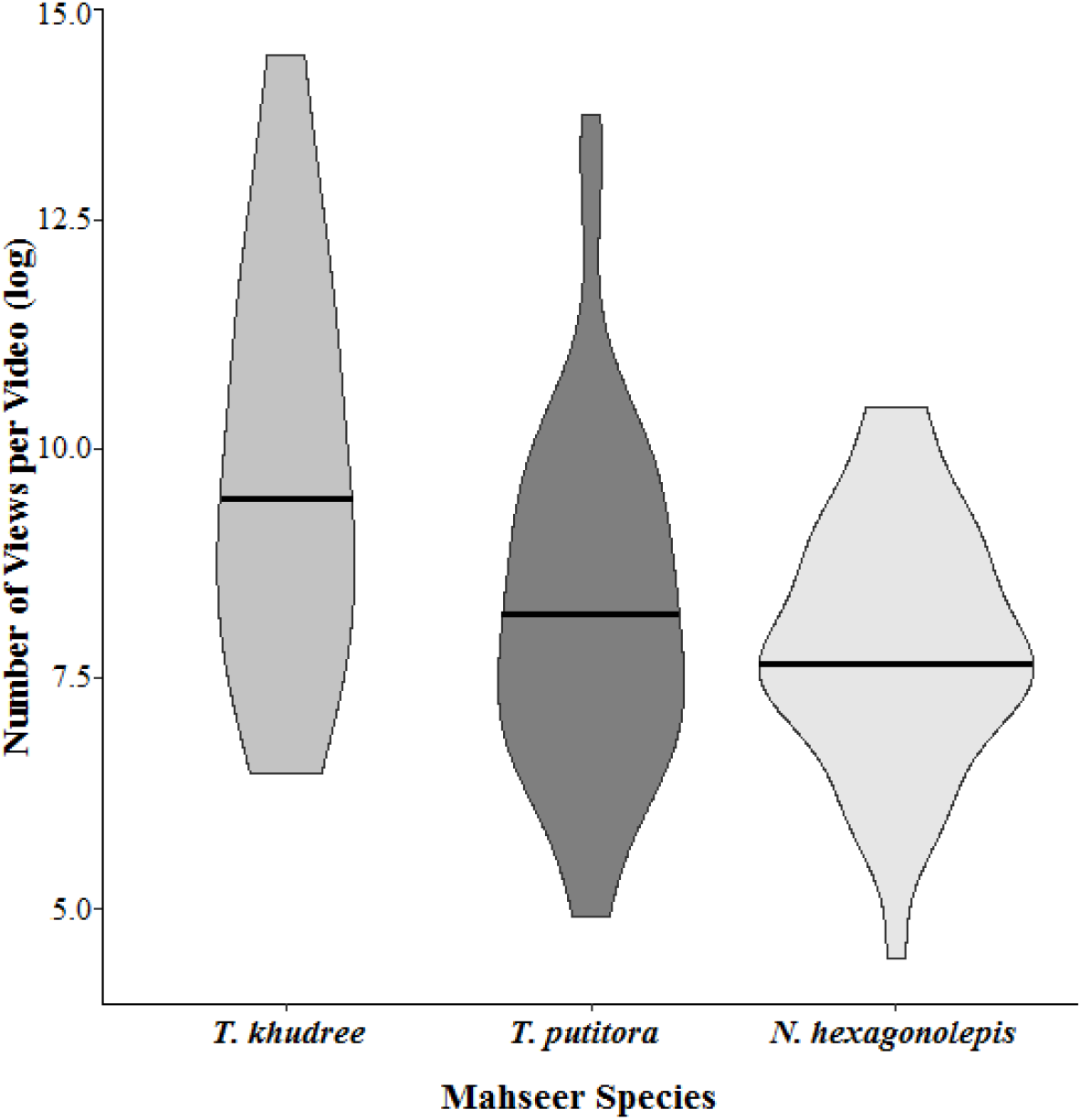
Number of views / video received by the angling videos of the three focal mahseer species on YouTube. The horizontal line across the plot represents the median.

**Figure 6.**
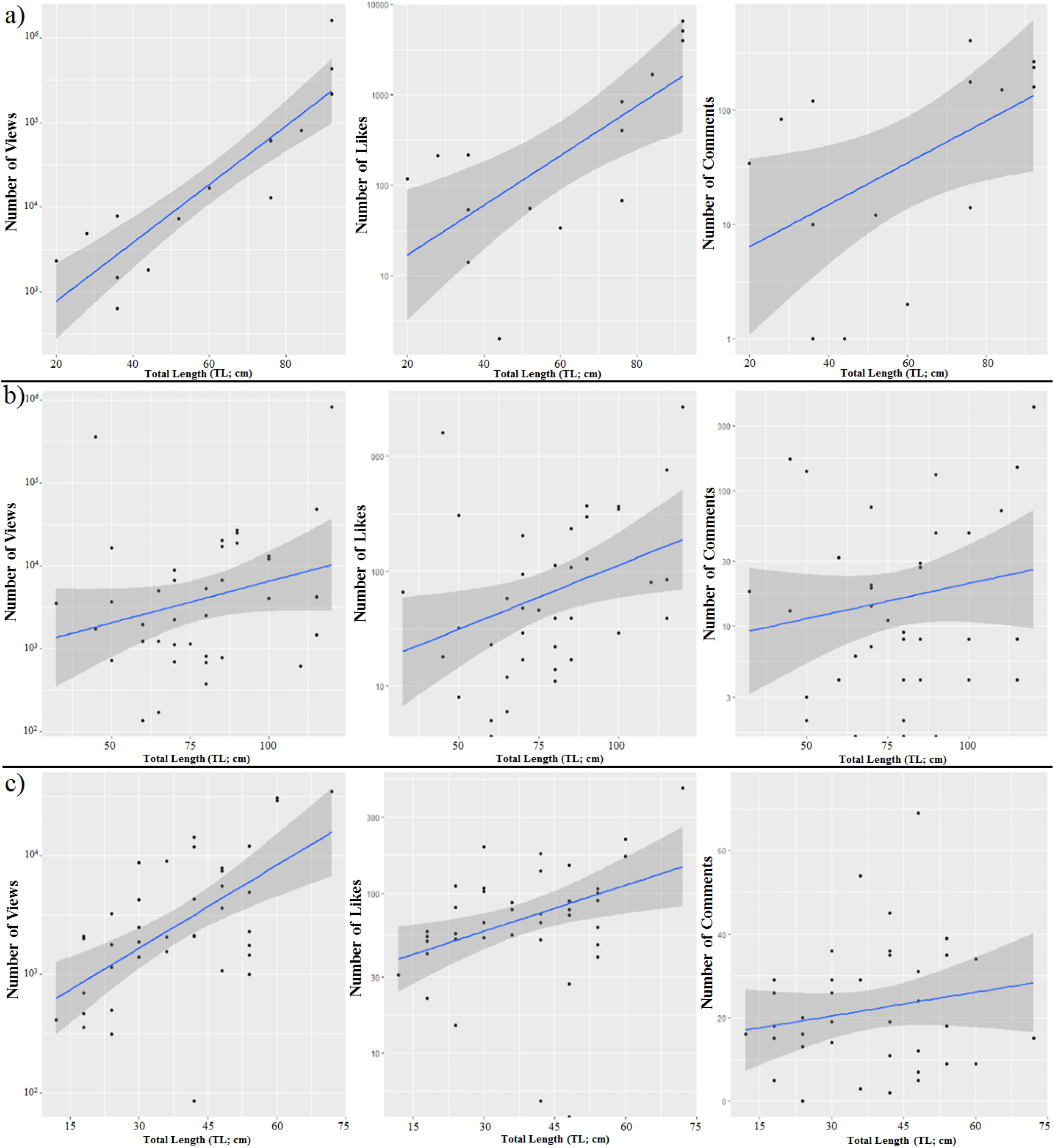
Relationship between the size (Total Length; TL) of the catch and Social Engagement (SE, number of views, likes, comments) received by the angling videos of a) *T. khudree,* b) *T. putitora* and c) *N. hexagonolepis*.

Dominant portions of the comments received by the angling videos of the three focal species were ‘angler related’ (*T. khudree* - 43.6%, *T. putitora -* 28.5% and *N. hexagonolepis -* 29.7%; Table 1). However, in the case of *N. hexagonolepis*, a lot of comments (29.2%) associated with ‘angling technique’ were also present. Interestingly, comments mentioning the IAF tilapia (1.2%) were only observed below the videos of *T. khudree*. The sentiment analysis of the comments using AFINN Lexicon resulted in a score of ‘+2’ for *T. khudree* and *T. putitora*, and ‘+3’ for *N. hexagonolepis* angling videos. We also checked the association between the sentiment scores and the size of the catch (TL); only in the case of *T. khudree* a significant association (positive) between these two variables were observed (β = 0.05, p = 0.02; Fig. 7). The emotional classification analysis revealed a higher prominence of words associated with the emotion ‘trust’ in the comments on *T. khudree*, *T. putitora* and *N. hexagonolepis* angling videos (between 25 - 35%; Fig. 8). The angler appreciation words ‘awesome’, ‘nice’, ‘good’, ‘great’ were the main keywords that emerged from the word frequency analysis of the comments on *T. khudree*, while both *T. putitora* and *N. hexagonolepis* videos showed more angling techniques and location related keywords (Table 2). The sentiment analysis of the comments received by the 15 videos which did not show any mahseer getting caught also revealed positive scores (+1 for *T. khudree* and +2 for both *T. putitora* and *N. hexagonolepis*).

**Figure 7.**
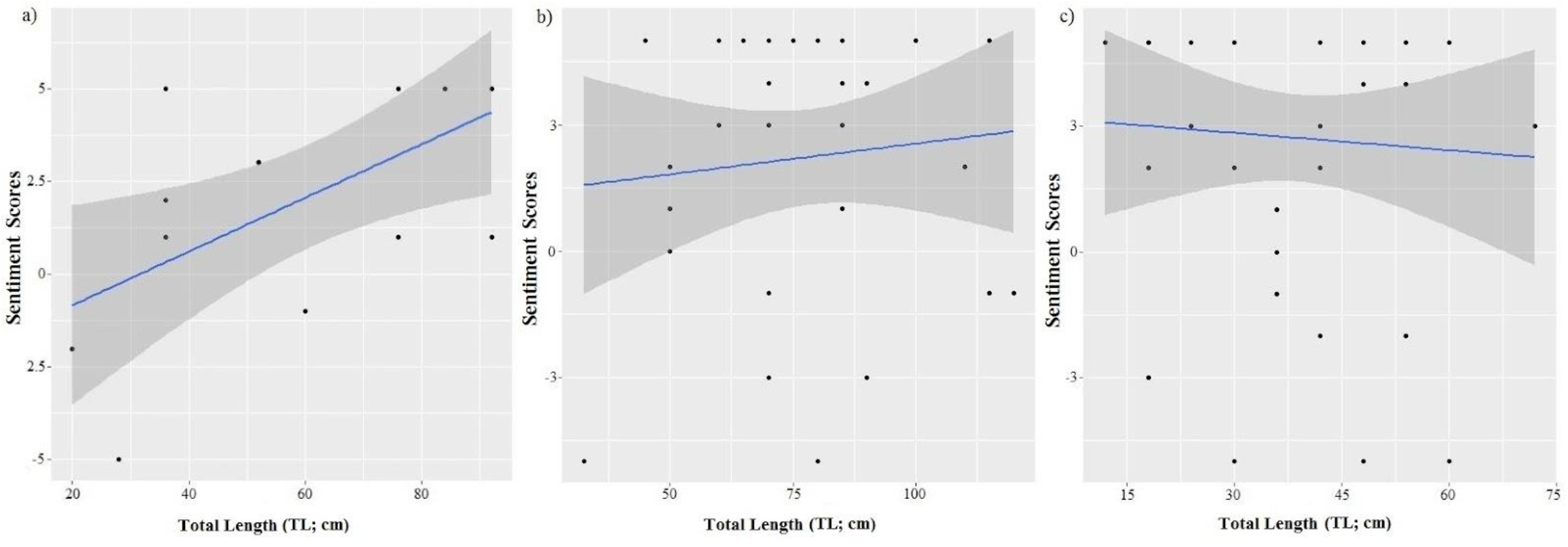
Relationship between the size (Total Length; TL) of the catch and the sentiment scores of the comments received on the angling videos of a) *T. khudree,* b) *T. putitora* and c) *N. hexagonolepis*.

**Figure 8.**
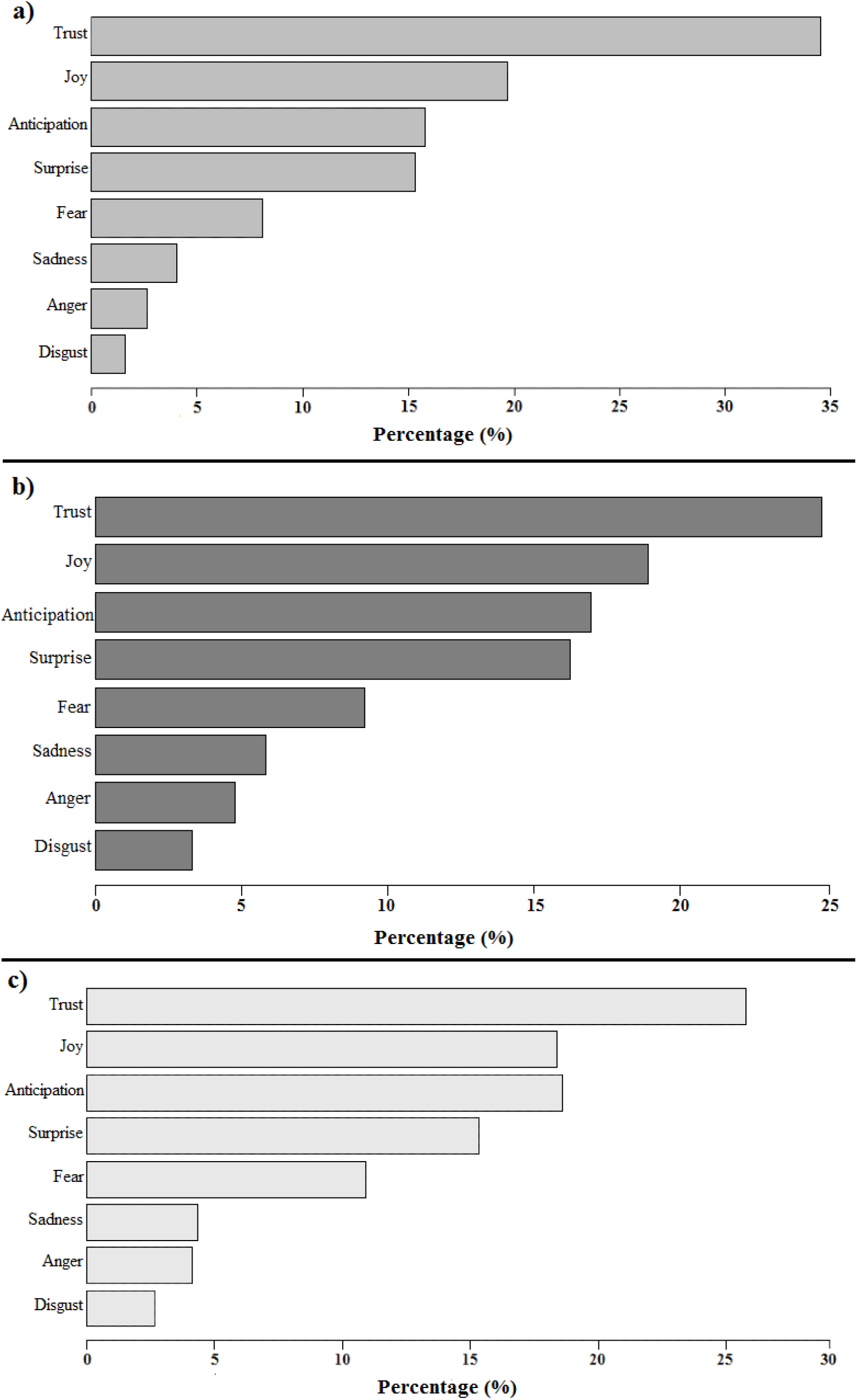
Emotional contents of the comments received by the angling videos of a) *T. khudree*, b) *T. putitora* and c) *N. hexagonolepis*.

**Table 1.** Details of the comments received by the angling videos of the three focal mahseer species.

| Category of Comments | Percentage (%) of Occurrence |  |  |
| --- | --- | --- | --- |
|  | *T. khudree* | *T. putitora* | *N. hexagonolepis* |
| Fish Related | 11.2 | 12.5 | 10.1 |
| Angler Related | 43.6 | 28.5 | 29.7 |
| Angling Technique Related | 15.0 | 21.0 | 29.2 |
| Location Related | 11.0 | 18.0 | 17.0 |
| Invasive Alien Fishes (IAF) Related | 1.2 | 0 | 0 |
| Miscellaneous | 18.0 | 20.0 | 14.0 |

**Table 2.**
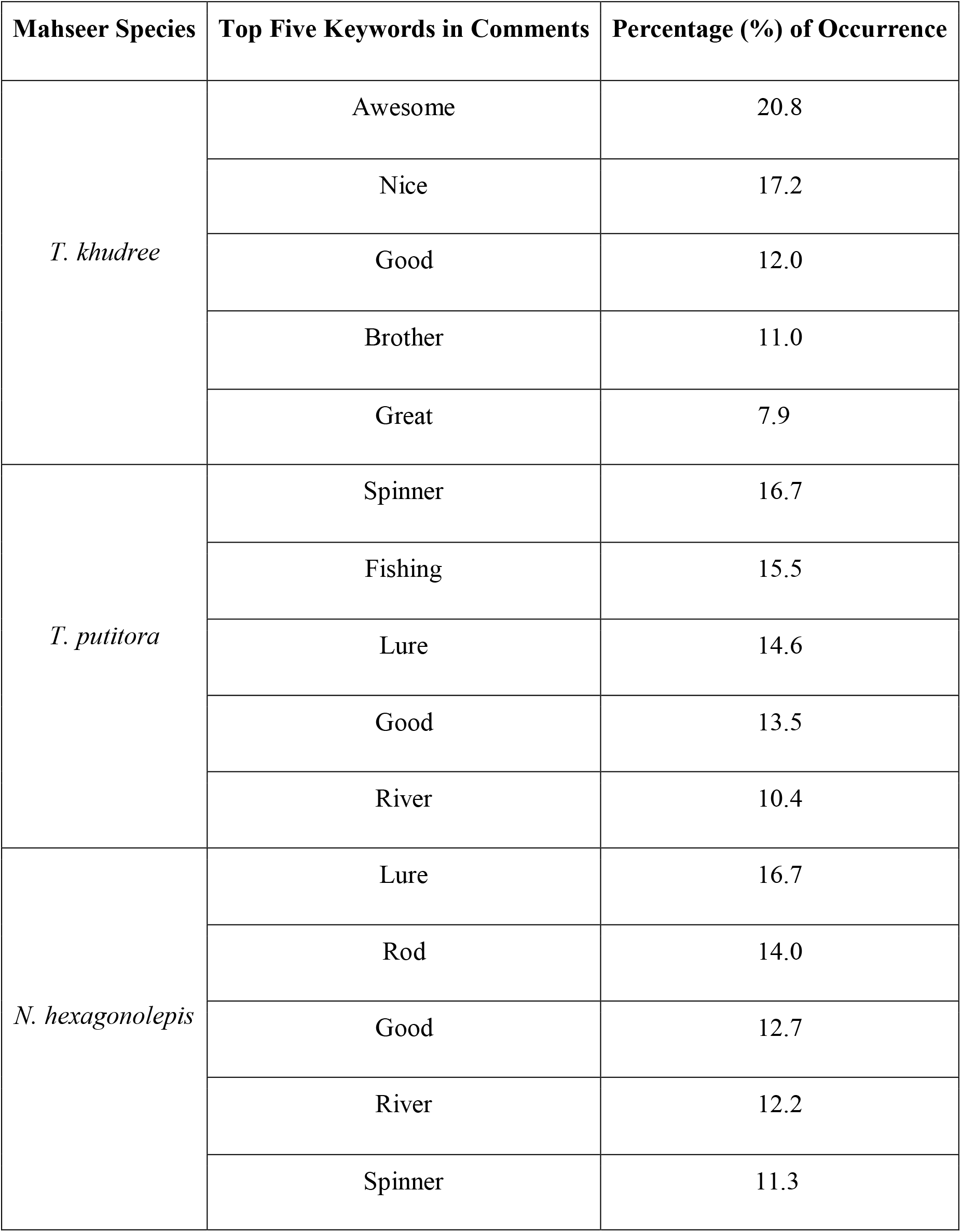
Frequency of occurrence of the top five keywords in the comments received by the angling videos of the three focal mahseer species.

| **Mahseer Species** | **Top Five Keywords in Comments** | **Percentage (%) of Occurrence** |
| --- | --- | --- |
| *T. khudree* | Awesome | 20.8 |
|  | Nice | 17.2 |
|  | Good | 12.0 |
|  | Brother | 11.0 |
|  | Great | 7.9 |
| *T. putitora* | Spinner | 16.7 |
|  | Fishing | 15.5 |
|  | Lure | 14.6 |
|  | Good | 13.5 |
|  | River | 10.4 |
| *N. hexagonolepis* | Lure | 16.7 |
|  | Rod | 14.0 |
|  | Good | 12.7 |
|  | River | 12.2 |
|  | Spinner | 11.3 |

## Discussion

Although using internet data as a complementary tool to get insights on various dimensions of RA is acquiring popularity amongst conservation scientists (Sbragaglia et al. 2020; Eryasar and Saygu 2022) and many anglers across the globe are uploading videos of their angling activities on social media such as YouTube, only 93 videos of mahseer angling activities undertaken in India in the last decade were available on YouTube. In comparison to the angling videos of marine fishes being added to this social media platform from different nations (Belhabib et al. 2016; Giovos et al. 2018; Sbragaglia et al. 2020; 2021; 2022; Eryasar and Saygu 2022), this number is noticeably small. Interestingly, even though during the time period 2010 - 2012, the internet, cheap mobile cameras and video editing tools were not much popular in India, a handful of angling videos of *T. khudree* and *T. putitora* were uploaded on YouTube. Later explorations revealed that these were mostly made as a part of tourism promotion and international sportfishing video documentation. Interestingly, no videos of mahseer angling conducted in India were uploaded on YouTube in the period from 2012 to 2015. Although tracing out the exact reason behind this unique observation requires further research, an amendment in the Indian Wildlife Protection Act (WPA 1972) by the Supreme Court of India in 2012 that banned angling from protected areas (Gupta et al. 2015b) might be a cause.

In India, many regions conduct mahseer angling festivals during the pre and post-monsoon months (Pinder et al. 2015; Baruah 2018; Baruah and Sarma 2018). However, no significant seasonal variation was observed in the number of angling videos of all three focal mahseer species uploaded on YouTube. This lack of seasonality in the upload frequency - assuming that the videos are uploaded within a few days of the angling activities (Sbragaglia et al. 2020) - points that mahseer species, which are intermittent breeders and batch spawners, laying eggs in multiple intervals (Khan 1939; Bhatnagar 1964; Sunder and Joshi 1976; Badola and Singh 1984; Sarma et al. 2018; Zaidi and Sarma DCFR n.d.), are angled throughout the year. This finding is of concern since the angling of breeders could negatively affect their lifespan (Lennox et al. 2022). Understanding the reproductive status of the angled individuals from the YouTube videos was not possible since many studies have reported heterogeneity in the body length at sexual maturity in different species of mahseers studied from various regions of India (Nautiyal and Lal 1985; Bhatt et al. 1998; Johal et al. 2000; Johal and Negi 2003; Mahapatra and Vinod 2011; Bhatt and Pandit 2016). Hence the anglers should be made aware to avoid further angling in areas where gravid individuals occur and they should be trained to handle the sexually mature fish caught on the hook properly to avoid injury and follow the catch and release protocol. RA can otherwise take a toll on the mahseer population in the near future.

None of the angling videos considered in the present study were recorded from locations outside the geographical distribution range of the focal mahseer species. In India, attempts were made to introduce popular mahseer species in the rivers outside the areas of their endemic distribution (Jha et al. 2018) in the past. Another concern raised by the present study is the presence of IAF tilapia, as ‘by-catch’ in *T. khudree* angling videos (3 in numbers) shot from the states of Maharashtra and Kerala. The IAF have evolved as a threat to the indigenous species globally and the local mahseer populations are not an exception to them (Raghavan et al. 2011; Gupta et al. 2020). We feel that these two results are incomplete and need further field-based and internet-based studies considering other social media platforms since only a fraction of mahseer angling activities would have been recorded and uploaded in the past due to the lack of availability of smartphones and the internet for the non-professional anglers in India. Furthermore, the presence of IAF in the mahseer videos should be taken seriously and attempts should be made to manage their populations in the natural water bodies before they become a menace to the indigenous mahseers (Raghavan et al. 2011; Gupta et al. 2020).

We did not come across any videos in which *T. mosal* and *T. remadevii* were angled. There is an unsolved confusion existing among researchers about the taxonomic position and distribution of *T. mosal,* a Data Deficient (DD) species in the IUCN Red List (Dahanukar et al. 2018) lacking voucher specimens. Making the situation further complex, in many contexts *T. mosal* has been used as a synonym for *T. putitora* (Menon 1999). Interestingly, neither the titles nor the descriptions of any mahseer angling videos studied mentioned the scientific or vernacular names of *T. mosal* or *T. remadevii*. Attempts to identify *T. mosal* and *T. remadevii* individuals amongst the mahseers portrayed in the angling videos using morphology-based keys (MacDonald 1948; Kurup and Radhakrishnan 2007; Nautiyal 2014; Pinder et al. 2018) also failed to give any positive results. No uploads of *T. remadevii,* the only *Tor* species present under the Critically Endangered category of the IUCN Red List (Pinder et al. 2018) may be due to their minimal presence in the freshwater habitats of India (Pinder et al. 2015a). A decline in their population dates back to the mid-2000s when catches gradually decreased (Pinder et al. 2015b). Additionally, no recent study evaluating their population size and distribution is available in the literature. Moreover, the name *T. remadevii,* got popularity in 2018 (Bangalore Mirror Bureau 2018; Sudhi 2018) though this species was first described in 2007 (Kurup and Radhakrishnan 2007).

The number of angling videos as well as the number of individuals hooked was the highest for *N. hexagonolepis* in comparison to the other species. The angling videos of *N. hexagonolepis* began to appear on YouTube in 2017 and attained a peak in 2022. Even the lockdown and travel restrictions implemented to mitigate the COVID-19 pandemic did not affect the number of *N. hexagonolepis* videos uploaded. Unfortunately, although many organisations in India have been promoting ethical angling, very few anglers were found to be practising C&R in the case of *N. hexagonolepis* (only 39% of individuals hooked were released). This species of mahseer is the most sought-after and highly-priced food fish with very high consumer demand in North-East India (Mahapatra et al 2004; Barua and Sarma 2018; Sikkim Government Gazette 2021). Furthermore, many indigenous Tribal and Nepali communities have been fishing for this ‘delicacy’ for generations (Baruah and Sarma 2018). Hence *N. hexagonolepis* may be angled in greater numbers by amateurs and local people in addition to professionals. Popularity and public awareness generated by the recognition of *N. hexagonolepis* as the ‘State Fish’ by the Indian state Sikkim in September 2021 (Sikkim Government Gazette 2021) could also have contributed to the increment in the number of videos of this species being uploaded. Our result is an alarm call to make the anglers aware of the ‘Near Threatened’ status of this species (Arunachalam 2010) and the need for following sustainable, responsible and ethical angling practices such as C&R to prevent depletion of their natural populations. A similar program may be required for the ‘endangered’ species, *T. putitora* as our result revealed that more than half (52.6%) of the individuals caught, were not released back into the rivers.

In all the three species of mahseers studied; the bigger the fish, the higher the Social Engagement (SE) it received. Previous studies focusing on terrestrial hunting (Child and Darimont 2015) and spearfishing (Sbragaglia et al. 2020) revealed a similar positive association between the SE and emotional signals with the catch size. Catching a bigger mahseer may be seen by the public as a ‘tough job done well’ and therefore trigger appreciation for the masculinity of the angler (Bull 2009), which in turn translates into a higher number of views, likes and comments (Child and Darimont 2015). The comments received by the angling videos of all three focal species of mahseer were related to anglers, angling locations or techniques, and included a significantly high proportion of positive mentions. The emotion that dominated these comments, ‘trust’ also indicates the support of viewers to the anglers. Although knowing the details of these YouTube video viewers is not possible, it seems that the main consumers of the mahseer angling videos are anglers or angling enthusiasts. The positive sentiment score obtained by the videos where no fish was caught but contained the term ‘mahseer angling’ in the title or the description also supports this argument. The viewers might have been able to understand the disappointment of a failed angler and therefore may not have discouraged them by giving negative comments or emoticons.

Amongst the three different parameters considered under the SE, only the number of views exhibited a species specific variation. *T. khudree* attracted a significantly large number of views per video and revealed a positive relation between the size of the catch (TL) and sentiment scores of the comments. Interestingly, a major portion of the angled *T. khudree* individuals (59.1%) were also found to be released back into their habitat. This higher viewership and positive comments obtained by the *T. khudree* videos could be taken as an indicator of people’s familiarity with and interest in this species. From the mid-1970, various organisations such as the Wildlife Association of South India (WASI n.d.), Coorg Wildlife Society (CWS n.d.), etc. have been conducting activities to enhance awareness, promote responsible and ethical C&R angling, and ensure the protection of this species. However, this increased viewership and SE attracted by the videos of *T. khudree* and larger individuals of *T. putitora* and *N. hexagonolepis* may persuade anglers to target these fishes. There is a high chance that anglers will target fishes which could provide them with increased rewards in the real and digital worlds. Additionally, in many contexts increased attention received on social media could also be translated into financial benefits (Han 2020). However, anglers should be made aware of the fact that larger *T. khudree* individuals, which may engage in longer fights, can undergo greater physiological impacts. The increased duration of air exposure, handling time and multiple angling episodes could result in reflex impairment and reduced movement over a period of 24 hours post release, increasing the susceptibility of post release mortality in larger individuals (Bower et al. 2016).

Although our study keeping the YouTube videos in focus gives many insights into various aspects of mahseer angling in India, we need to acknowledge the following limitations; lack of availability of large numbers of videos and focus on a single social media platform. In India along with YouTube; Twitter, Facebook and Instagram are also very popular (Kemp 2023; OOSGA 2023) and these platforms also provide the facility to upload videos. Hence, to get an elucidated picture of mahseer angling in India it is essential to conduct an analysis integrating the field-based studies focusing on the behaviour of anglers, perceptions and attitudes of various stakeholders towards multiple mahseer species and the different materials on mahseer RA being uploaded on the internet and various social media channels. The results of such a study would function as a catalyst to develop an internet data-based complementary tool to understand and monitor the mahseer RA in India and may provide significant insights to develop, implement and refine plans for protecting the natural populations of these valuable megafishes.

## Conclusion

In India, RA has been propagated as a tool to protect mahseer populations in the wild by many organisations (Everard and Kataria 2011; Pinder and Raghavan 2013; Baruah and Sarma 2018). This model aims to generate monetary benefits and livelihood for the local people from RA and associated tourism activities, hence motivating them to protect mahseer from poaching and other forms of illegal fishing activities (Everard et al. 2009). This plan had been implemented in different riverine ecosystems in India such as Bhikiyasen - Ramganga River Uttarakhand (Everard & Kataria 2011), Kodagu - Cauvery River, Karnataka (Pinder and Raghavan 2013), Nameri - Jia Bhorelli River, Assam (Barauh and Sarma 2018), etc. However, in a geographically large nation like India, it is not easy to monitor the angling activities and suggest modifications in practices and implement evidence based policy reformations to ensure the well-being of the mahseer populations being exploited for RA. In this context, studies like the present one focusing on social media and other internet data could work as a complementary mechanism to get insights into the angling activities and the public response towards them. Furthermore, the study of angling videos could also help to understand the presence and extension of the invasive alien fish species (IAF) in natural water bodies. Although currently in its infancy in India, future conservation culturomics and iEcology studies focusing on the highly exploited and vulnerable populations of megafishes like mahseer, are expected to evolve as complementary tools for scientists, policymakers and ecosystem managers working to protect the freshwater fish diversity.

## Acknowledgements

Prantik Das acknowledges the Council of Scientific and Industrial Research (CSIR) - Human Resource Development Group (HRDG), New Delhi for the research fellowship. Binoy expresses gratitude to the Science and Engineering Research Board, India (SERB-CRG/2021/007778).

## Supplementary Materials

**SM1:**
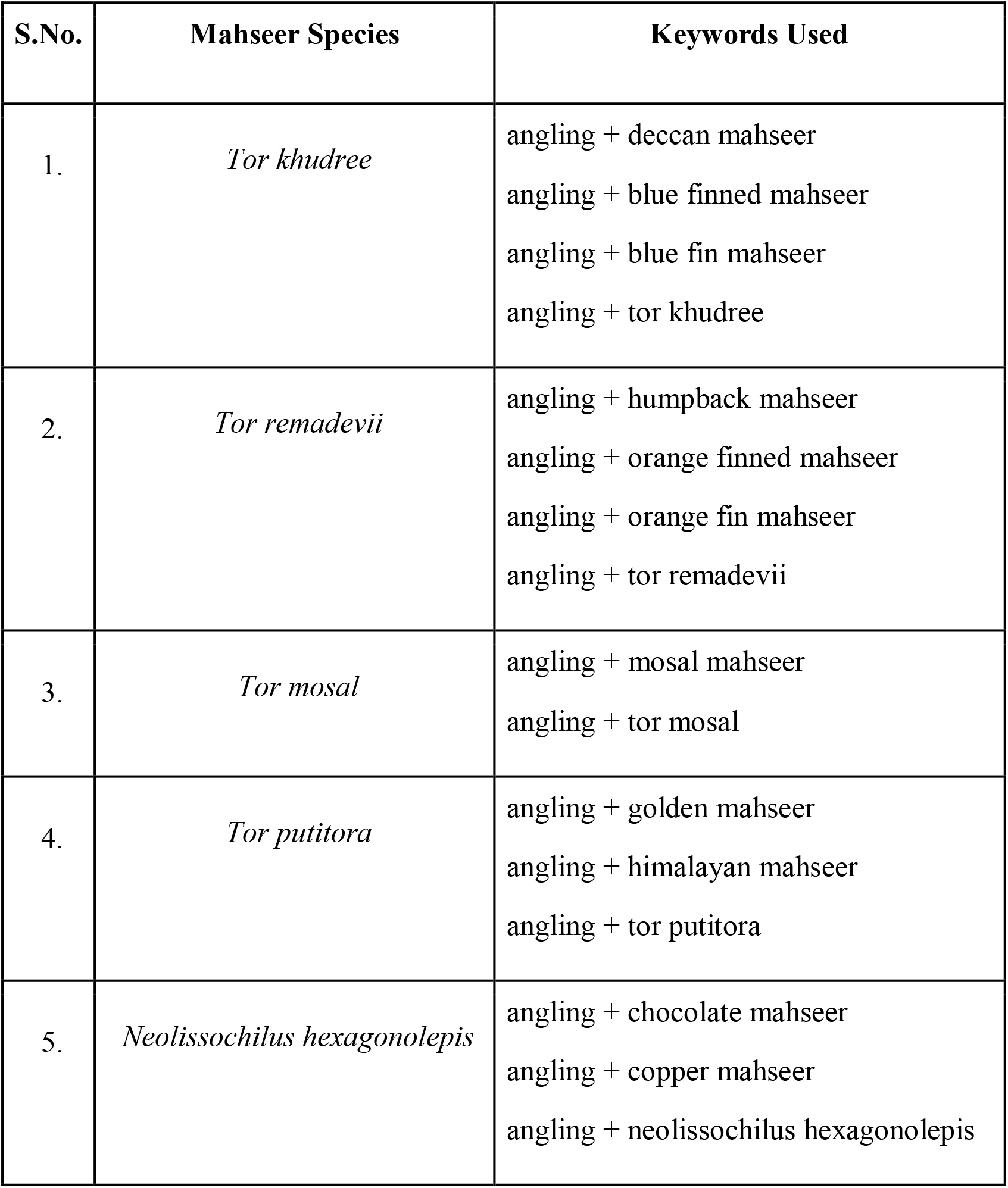
List of the keywords used for searching videos of the five mahseer species popularly angled in India, on YouTube.

