## Supplemental Material (SM1) for "Landing the ‘Tiger of Rivers’: Understanding Recreational Angling of Mahseers in India using YouTube Videos"

**Supplementary Materials**

**SM1:** List of the keywords used for searching videos of the five mahseer species popularly angled in India, on YouTube.

| **S.No.** | **Mahseer Species** | **Keywords Used** |
| --- | --- | --- |
| 1. | *Tor khudree* | angling + deccan mahseer  angling + blue finned mahseer  angling + blue fin mahseer  angling + tor khudree |
| 2. | *Tor remadevii* | angling + humpback mahseer  angling + orange finned mahseer  angling + orange fin mahseer  angling + tor remadevii |
| 3. | *Tor mosal* | angling + mosal mahseer  angling + tor mosal |
| 4. | *Tor putitora* | angling + golden mahseer  angling + himalayan mahseer  angling + tor putitora |
| 5. | *Neolissochilus hexagonolepis* | angling + chocolate mahseer  angling + copper mahseer  angling + neolissochilus hexagonolepis |

_____________________
